# Yeast–Plant-Derived Consortia Factors Exhibit A Distinct And Enriched Beneficial Metabolome For Health And Beauty

**DOI:** 10.64898/2026.01.29.702615

**Authors:** Assaf Zemach, Mikhail R. Plaza, Dior Simmons, Bong Seop Lee, Melissa Palomares, Dodanim Talavera-Adame, Nathan Newman

## Abstract

Cells secrete metabolites and other factors into the extracellular medium, collectively referred to as the secretome. The effect of co-culturing cells from different species on their combined secretome remains poorly understood. Here, we investigated the effect of co-culturing plant and yeast cells to produce a unique set of secretory factors collectively termed Consortia Factors (CFx). Specifically, a yeast suspension of *Ustilago cynodontis* (Ustilago) was co-cultured with plant cells derived from *Ocimum sanctum* (Tulsi). The metabolome of the Ustilago-Tulsi CFx (termed CFx-α1) was then profiled and compared with that of the individual cultures. Statistical clustering analyses revealed that the CFx-α1 metabolome was substantially different from either culture grown alone and was enriched in metabolites relative to single cultures. Notably, significant enrichment was observed among metabolites mutually upregulated in CFx compared with both single cultures, with approximately one-third absent from either culture alone. Using an algorithm designed to identify scientifically validated antioxidant metabolites, the CFx-α1 was found to be enriched in antioxidants relative to the single cultures, including vitamin C. Accordingly, the CFx-α1 exhibited stronger antioxidant activity than either of the single-culture secretomes, and even than the combination of the two single-culture secretomes. These findings suggest that consortia factors derived from co-culturing yeast and plant cells can generate a unique and more potent secretome that is enriched in bioactive metabolites beneficial for human health.

## Introduction

The secretome is the complete set of biochemical factors secreted by cells into the extracellular space (1). It plays essential roles in numerous biological processes and interactions (2). A significant component of the secretome is metabolites—low-molecular-weight molecules that function as key intermediates and products of metabolism. These include sugars, amino acids, lipids, nucleotides, organic acids, vitamins, hormones, flavonoids, phenolics, and glucosinolates. Advances in analytical technologies have greatly improved the detection and quantification of these metabolites, enabling detailed investigation of metabolic networks and cellular physiology (3).

Plants and fungi synthesize a wide range of metabolites that contribute significantly to human health. Yeast, for example, produces both primary and secondary metabolites with major industrial relevance. These include biofuels such as bioethanol, organic acids, hydrolytic enzymes, vitamins, amino acids, and antibiotics-products vital to the food, agricultural, and pharmaceutical sectors (4). Plant cell suspension cultures have also emerged as a valuable biotechnological platform for producing high-value secondary metabolites used in pharmaceuticals, cosmetics, and food. Such cultures offer key advantages: controlled production conditions, independence from climate and soil, and year-round yields of consistent quality and quantity (5–7). The secretome of plant cell suspensions, which include extracellular proteins and metabolites, is central to these processes. Because it can be separated from cells without disruption, the secretome provides a continuous and reliable source of extracellular proteins essential for cell wall remodeling and stress defense (8).

The potential of interspecies cell cultures has been increasingly recognized. For instance, endophytic fungi inhabit plant tissues in symbiosis and produce diverse bioactive compounds with health benefits such as antioxidant, anti-inflammatory, antimicrobial, and anticancer activities (9). Similarly, microalgae–yeast co-cultivation creates a symbiotic system that produces valuable biofuels by sharing nutrients and metabolic byproducts, thereby enhancing overall lipid and biomass productivity for sustainable energy (10). These examples affirm the benefits of interspecies co-culturing for metabolite production.

The Tulsi plant (*Ocimum tenuiflorum/sanctum*), also known as Holy basil, is a revered herb in traditional medicine, particularly in Ayurveda. Its leaves contain diverse compounds—including phenolics, flavonoids, and terpenoids—that contribute to pharmacological activities such as anti-stress, anticancer, antioxidant, and antidiabetic effects (11,12). The fungus *Ustilago cynodontis*, a member of the Ustilaginaceae family, is typically known as a plant pathogen but also holds biotechnological importance (13,14). It produces bioactive compounds such as the organic acid itaconate and mannosylerythritol lipids, which have applications in cosmetics, polymers, and biosurfactants (15,16). Given the metabolic diversity observed in related Ustilaginaceae species (17), *U. cynodontis* remains a promising source for the discovery of additional bioactive compounds. In this study, untargeted metabolomics was applied to analyze and quantify the global effects of the secretome derived from an integrated cell suspension of the fungus *U. cynodontis* and the plant O. *sanctum*. The findings from the Ustilago–Tulsi co-culture, compared with either single culture or with a simple combination of their respective secretome, demonstrate that CFx-α1 generated by this patented process(18) yield a unique and significantly enriched repertoire of beneficial bioactive compounds.

## Materials and Methods

### Biological materials

Tulsi stem cell culture (callus) was established from an O. *sanctum* plant using plant growth hormones. The fungus *U. cynodontis* used in this study was isolated from a local bermudagrass *(Cynodon dactylon)* plant and was identified by its Internal Transcribed Spacer (ITS) sequence.*U. cynodontis* was cultivated in its yeast life cycle phase using an in-house-developed growth medium. ITS and whole-genome sequencing verified the identity of the fungus as *U. cynodontis*.

### Growth conditions

Ustilago and Tulsi cell cultures were cultivated in flasks containing Murashige and Skoog (MS)-based liquid medium supplemented with 3% sucrose and plant growth hormones (2,4-dichlorophenoxyacetic acid and kinetin). All cultures contained 50 mL of medium. Tulsi cultures were inoculated with 5 g of callus, Ustilago cultures with 4 × 107 cells, and Ustilago–Tulsi co-cultures with 5 g of Tulsi cells and 4 × 107 Ustilago cells (18). Flasks were shaken at 120 rpm on an orbital shaker at room temperature (25 °C ± 1 °C). Growth was measured by biomass fresh weight and sugar consumption. Following seven days of cultivation, culture media were separated by centrifugation and stored at −80 °C. Three biological replicates were generated for each of the cultures: Tulsi, Ustilago, and Ustilago–Tulsi.

### Ustilago genome sequencing

DNA extraction was performed on *U. cynodontis* cell pellets using the Genologue Universal Microbiome Extraction Kit (Genologue, Tucker, GA), following the manufacturer’s instructions. Library construction was carried out according to standard Illumina TruSeq chemistry using a modified KAPA HyperPrep kit and protocol. DNA was mechanically sheared to approximately 400 bp (peak size) by ultrasonication using a Bioruptor sonicator (Diagenode, Denville, NJ). Approximately 100 ng of fragmented DNA first underwent an end-repair reaction to form blunt ends, followed by an A-tailing reaction to add 3′-A overhangs. A shortened version of Y-shaped adapters was then ligated to the DNA ends. Following bead-based double-sided size selection (Genologue, Tucker, GA), the DNA libraries were amplified using primers to incorporate Unique Dual Indexes (UDI, Eurofins Genomics, Louisville, KY) at both ends. The resulting libraries were normalized, pooled, and sequenced on a NovaSeq 6000 sequencer (Illumina, San Diego, CA) for paired-end 150 bp reads.

The resulting genomic data was quality controlled to remove low quality reads and adapter sequences and then aligned to a reference *U. cynodontis* genome (19) using the BWA short-read aligner (20). Variant calling was performed using *bcftools* (21) to identify SNPs, insertions, and deletions. The program CNVkit (22) was used to determine any copy number variation between our strain and the reference genome. The BRAKER2 pipeline was used to annotate and predict genes using gene prediction tools GeneMark-ES and AUGUSTUS (23,24). To determine the effect of mutation, indels, or copy number variation (CNV) has on predicted genes, the program SnpEff (25) was used to predict their functional effect. Finally, a phylogenetic tree using SANS serif (26) was created to confirm our strain’s relatedness to the *U. cynodontis* reference genome.

### Metabolomic analysis

Media derived from the three biological cultures of Ustilago, Tulsi, and Ustilago–Tulsi were submitted for untargeted metabolomics at the Carver Metabolomics Core, University of Illinois Urbana–Champaign Roy J. Carver Biotechnology Center (Urbana, IL). Processed samples were dried, reconstituted in 100 µL methanol:water, and 5 µL was injected onto the instrument for analysis. Samples were analyzed using a Dionex Ultimate 3000 series UHPLC system (Thermo Scientific) coupled to a Q-Exactive MS system (Thermo Scientific), as described previously (27). Samples were analyzed by reversed-phase liquid chromatography (RPLC) using a Waters Acquity ethylene-bridged hybrid (BEH) C18 column (100 mm × 2.1 mm, 1.7 μm) maintained at 25 °C with a flow rate of 0.3–0.4 mL/min (Waters Corp). The mobile phases consisted of (A) water with 0.1% formic acid and (B) acetonitrile with 0.1% formic acid. Spectra were acquired in both positive and negative ionization modes.

All LC–MS raw data files were processed using MS-DIAL v4.90 for data collection, peak detection, alignment, adduct assignment, and compound identification (28). The detailed parameter settings were as follows: MS1 tolerance, 0.005 Da; MS2 tolerance, 0.01 Da; minimum peak height, 10,000 amplitude units; mass slice width, 0.05 Da; smoothing method, linear weighted moving average; smoothing level, 3 scans; minimum peak width, 5 scans. [M-H]-, [2M-H]- and [M+H]+, [2M+H]+, [M+NH4]+, [M+Na]+, [M+2H]2+ were included as adducts in negative and positive modes, respectively. Compounds were annotated by m/z, MS/MS spectra, and retention time against an in-house library generated from chemical standards. In addition, they were annotated by m/z and MS/MS spectra using the public libraries MassBank of North America (MoNA), MassBank Europe (MassBank EU), and Global Natural Products Social Molecular Networking (GNPS), as well as the commercial library National Institute of Technology 20 (NIST20). Internal standards were monitored for retention time and intensity, and PCA was used for multivariate statistics and visualization, specifically for outlier detection.

From the MS-DIAL results file, all detected features/metabolites were removed if (sample max)/(blank average) < 10. Known (positively identified/annotated) features/metabolites were evaluated for MSI level 1 matches (29). Next, positive and negative mode data were combined and replicate identifications removed by initially retaining features with an MSI level 1 match. Because MS-DIAL does not evaluate MS/MS data for metabolites with both m/z and retention time matches, manual evaluation of spectra was performed to confirm level 1 identifications. Manual inspection of spectra for MSI level 2 matches was conducted in specific cases flagged by parameters requiring further investigation. Remaining replicate features without an MSI level 1 match were filtered by retaining those with the highest MS-DIAL total score. Detected synthetic drugs were removed from the dataset because they were not relevant to metabolite changes in response to the experimental treatment. For all features not positively identified (unknowns), after removal based on the above sample max/blank average threshold, unknowns with both m/z and MS2 data were retained and are reported separately. All sample peak heights (semi-quantitative) were normalized to the metabolite total ion chromatogram (mTIC) for each analysis (RPLC–positive, RPLC–negative). To perform statistical analyses, processed peak intensity data above the 100,000 threshold for compound detection from the *U. cynodontis*, Tulsi callus, and co-culture samples were uploaded to MetaboAnalyst 6.0 (30). Peak intensity data were log-transformed, and MetaboAnalyst was used to generate PCA plots and heatmaps to describe unique metabolomic profiles among the sample groups.

### Identification of metabolites with antioxidative properties

Untargeted metabolomic analysis on Ustilago and plant callus samples identified 1478 known metabolites. These known metabolites were searched against the PubMed database to discover abstracts that support whether they have antioxidative properties. Specifically, we searched using the phrase ({metabolite}[TIAB] AND (antiox* OR anti-ox*)[TIAB]) to find abstracts that had both the metabolite and any antioxidant name derivative in either the title and/or abstract. Titles and abstracts were downloaded from papers that matched this search and subsequently filtered to remove non-matching metabolite entries. Specifically, abstracts were excluded if the text contained target metabolite name mismatches or partial matches. The filtered title and abstract text data were used as input fortext annotation using OpenAI’s large language model GPT-4.1 mini (ChatGPT). The ChatGPT answers were downloaded and a subset of annotations were cross verified manually for accuracy. GPT and manual annotation data were imported into R to determine how accurate ChatGPT was in annotating titles and abstracts for metabolites having antioxidant activity. To assess this model, we determined the probability that a certain number of “Yes” by GPT would correctly correspond to human annotations. Metabolites were grouped into categories based on the number of GPT “Yes” annotations. For each category, ranging from 1 to 10 “Yes” annotations, four metabolites were randomly sampled. Each sample dataset consisted of 220 papers representing 40 different metabolites.

For each “Yes” count threshold (1 – 10 “Yes”), the probability was calculated for correct identification by GPT. There had to be at least one “Yes” manual annotation that matched to a GPT “Yes” annotation to be labeled as correct. The probability was calculated as the ratio of correct identifications (metabolites with at least one “Yes” manual annotation) to the total number of selected metabolites. This ratio represents the likelihood that a GPT annotation is correct when it meets or exceeds the given threshold:

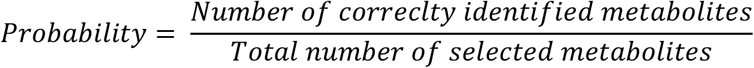

Where the number of correctly identified metabolites is greater or equal to one “Yes” manual annotation and the total number of selected metabolites the count of metabolites that is greater than or equal to the threshold. This process was then repeated for a total of 10,000 times creating different pools of metabolite and paper samples and the probability mean and standard deviation was calculated.

### 2,2-Diphenyl-1-picrylhydrazyl (DPPH) radical scavenging assay

The radical scavenging activity of culture media and standard compounds (vitamin C) was determined spectrophotometrically using the DPPH method (31). The reaction was set up by mixing 0.1 mL of test sample with 0.9 mL of DPPH (81 µM in methanol) and incubating the mixture in the dark at room temperature for 24 h. Absorbance was measured at 517 nm using a BioTek Synergy LX spectrophotometer. Measurements were performed on three biological replicates. Results were expressed in micromolar equivalents of vitamin C (L-ascorbic acid), which was calibrated using concentrations from 0 to 500 µM.

## Results

### Development and genome sequencing of Ustilago and Tulsi cell cultures

Tulsi cell culture was established from an O. sanctum plant (Figure 1A). *U. cynodontis* (Ustilago) was isolated from a local bermudagrass plant (Figure 1A). Species identification of the Ustilago culture was performed by ITS sequencing (Figure S1A–B). To further investigate the relationship of this Ustilago strain to other Ustilaginaceae species, its genome was sequenced and compared with 12 other Ustilaginaceae genomes. A phylogenetic tree based on genomic sequencing clustered this Ustilago strain together with the reference *U. cynodontis* genome, confirming its species identity (Figure 1B). Further analysis of the genome of this *U. cynodontis* strain identified 16,694 genetic variants relative to the published *U. cynodontis* genome (Figure S1C). Genetic variants were relatively enriched in repeat regions and depleted in gene-coding sequences (Figure S1C). In genic exons, the genetic variant frequency was approximately 3 per 10 kb, whereas in repeat sequences it was closer to 23 per 10 kb, about sevenfold higher (Figure S1C).

**Figure 1.**
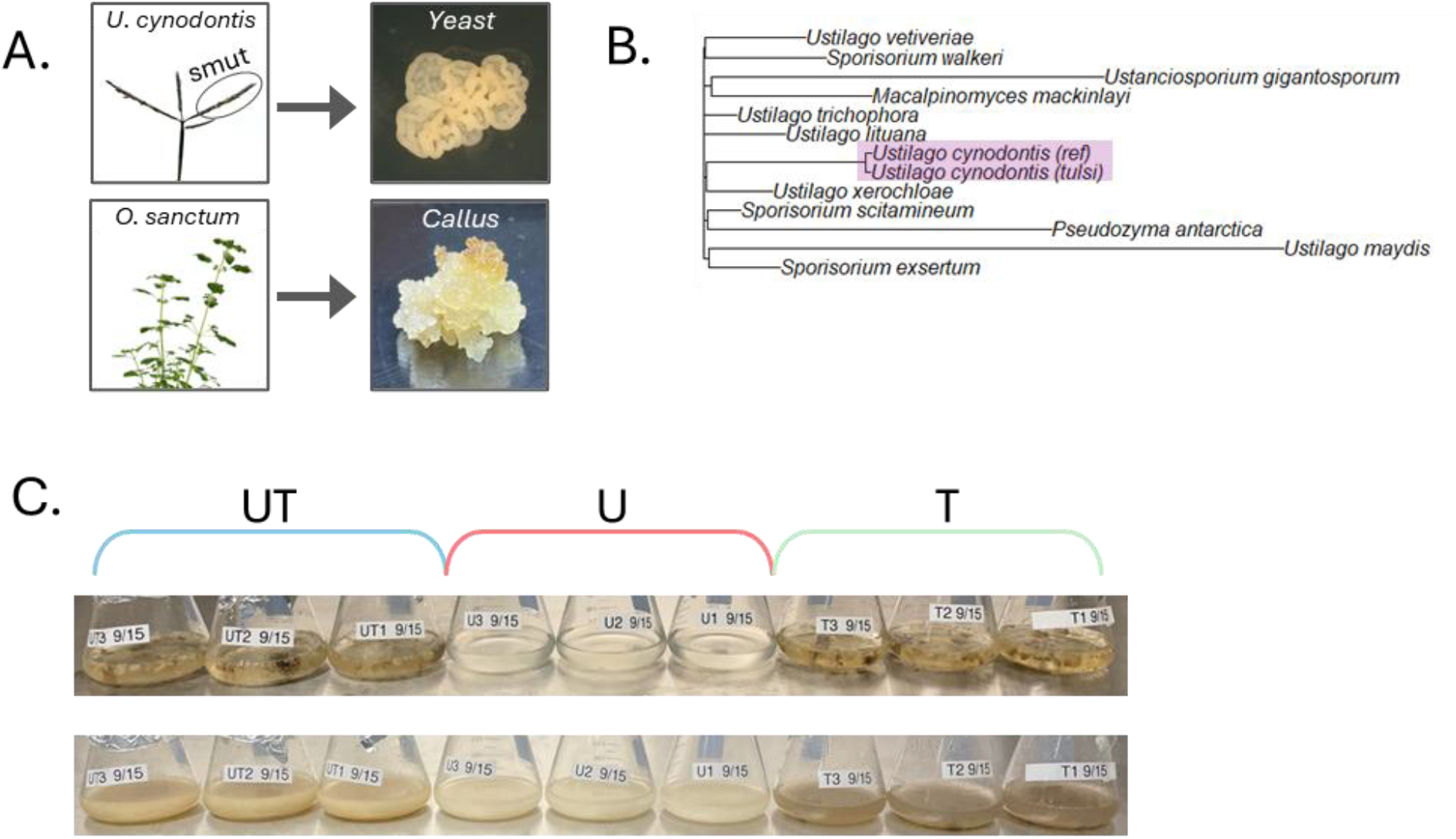
Co-culturing Ustilago (yeast) and Tulsi (plant) cells. A. Isolation of Ustilago cynodontis from a Bermudagrass plant (top), and the development of Tulsi cell culture (bottom). B. A phylogenetic tree based on the genomes of 12 Ustilaginaceae species and the one isolated in this study. C. Tulsi (T), Ustilago (U), and Ustilago-Tulsi (UT) Cultures at induction (top) and at harvest (bottom).

### CFx-α1 unique metabolome

Ustilago (U) and Tulsi (T) cells from callus were cultivated in flasks either separately or together (UT) (Figure 1C). Harvested cultures were centrifuged, and their corresponding media were subjected to untargeted metabolomics by LC–MS. Principal component analysis (PCA), hierarchical clustering, and Pearson correlation of the identified metabolite profiles separated the plant and yeast samples into distinct clusters (Figure 2A–C). The UT co-culture metabolomes, termed CFx-α1, formed a distinct cluster from either of the single cultures, while clustering more closely with the Ustilago samples than with the Tulsi samples (Figure 2A–C). In addition, the CFx-α1 metabolite profile was distinct from the simple sum of the single cultures (U + T), indicating that the CFx-α1 differs from a mere mixture of single-culture secretomes (Figure 2D). Overall, these results indicate that the CFx-α1 secretome is unique and distinct from the secretomes of the single cultures, either separately or mixed post-harvest.

**Figure 2.**
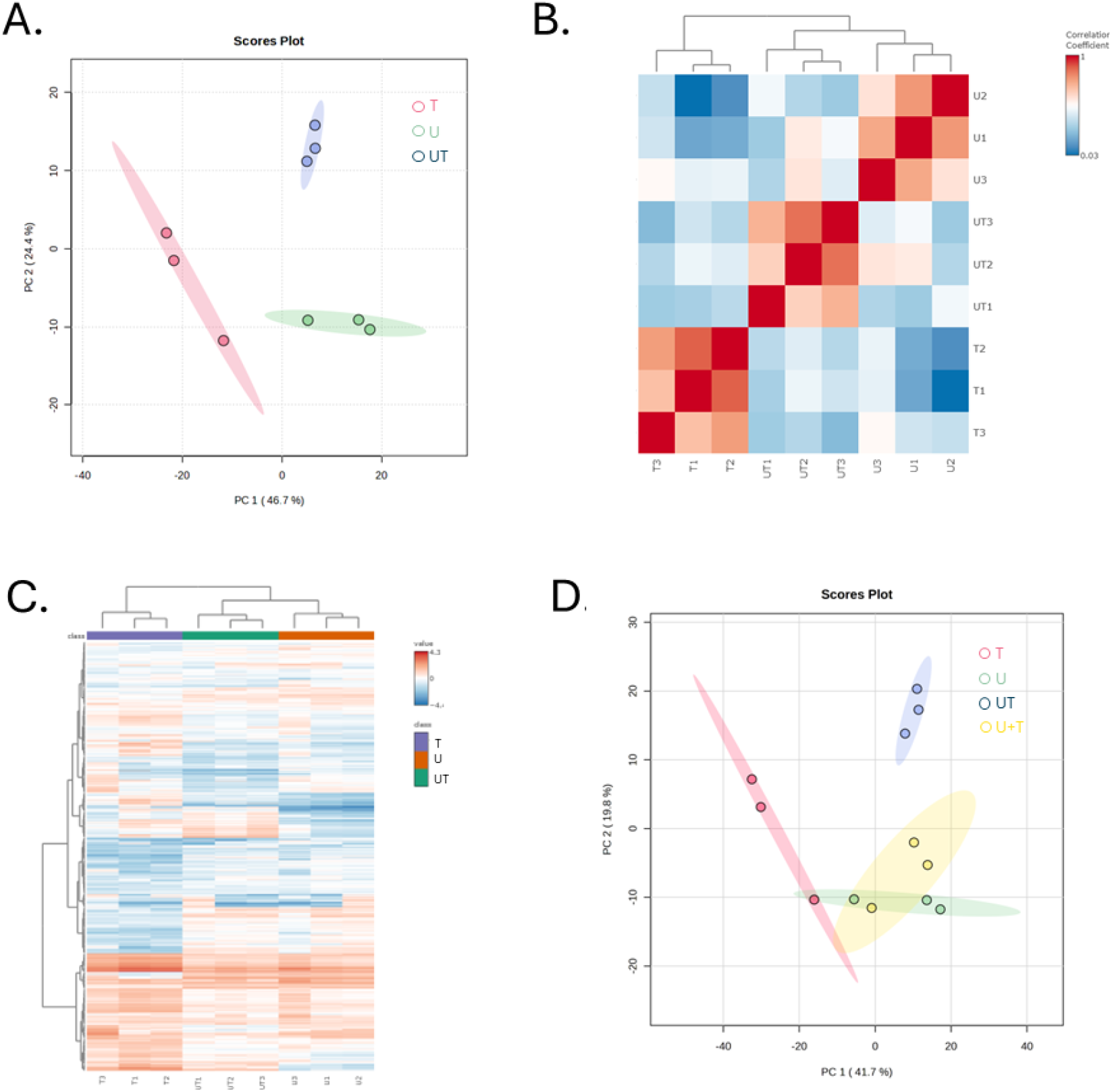
CFx-α1 unique metabolome. A-C. Clustering of LC-MS intensity (median normalized and log transformed) of identified metabolites (MSI1 & MSI2) in the three replicates of each of the cultures as principal component analysis (PCA) (A), Pearson correlation clustering (B), and Euclidean distance with Ward’s linkage hierarchical clustering (C). U, T, and UT stand for Ustilago, Tulsi, and Ustilago-Tulsi (CFx-α1), respectively. D. PCA as in A, including pseudo mixed samples of the single cultures, i.e., Ustilago plus Tulsi (U+T).

### CFx-α1 enriched metabolome

Next, metabolites that were differentially regulated in the CFx-α1 compared with the secretomes of U or T cultured individually were identified (Figure 3A). This analysis revealed that the CFx-α1 is relatively enriched in metabolites compared with secretomes derived from single cultures. Among the differentially regulated metabolites (DRMs), more metabolites were upregulated in CFx-α1 than in either of the single cultures (Figure 3B). CFx-α1 contained 432 and 163 upregulated DRMs (up-DRMs) relative to Tulsi and Ustilago, respectively, which is approximately fourfold and twofold higher than the corresponding numbers of downregulated DRMs (down-DRMs) in CFx-α1 versus T (98) and U (82). Additionally, the relative fold change of CFx-α1 up-DRMs was two-to threefold higher than that of down-DRMs (Figure 3B).

**Figure 3.**
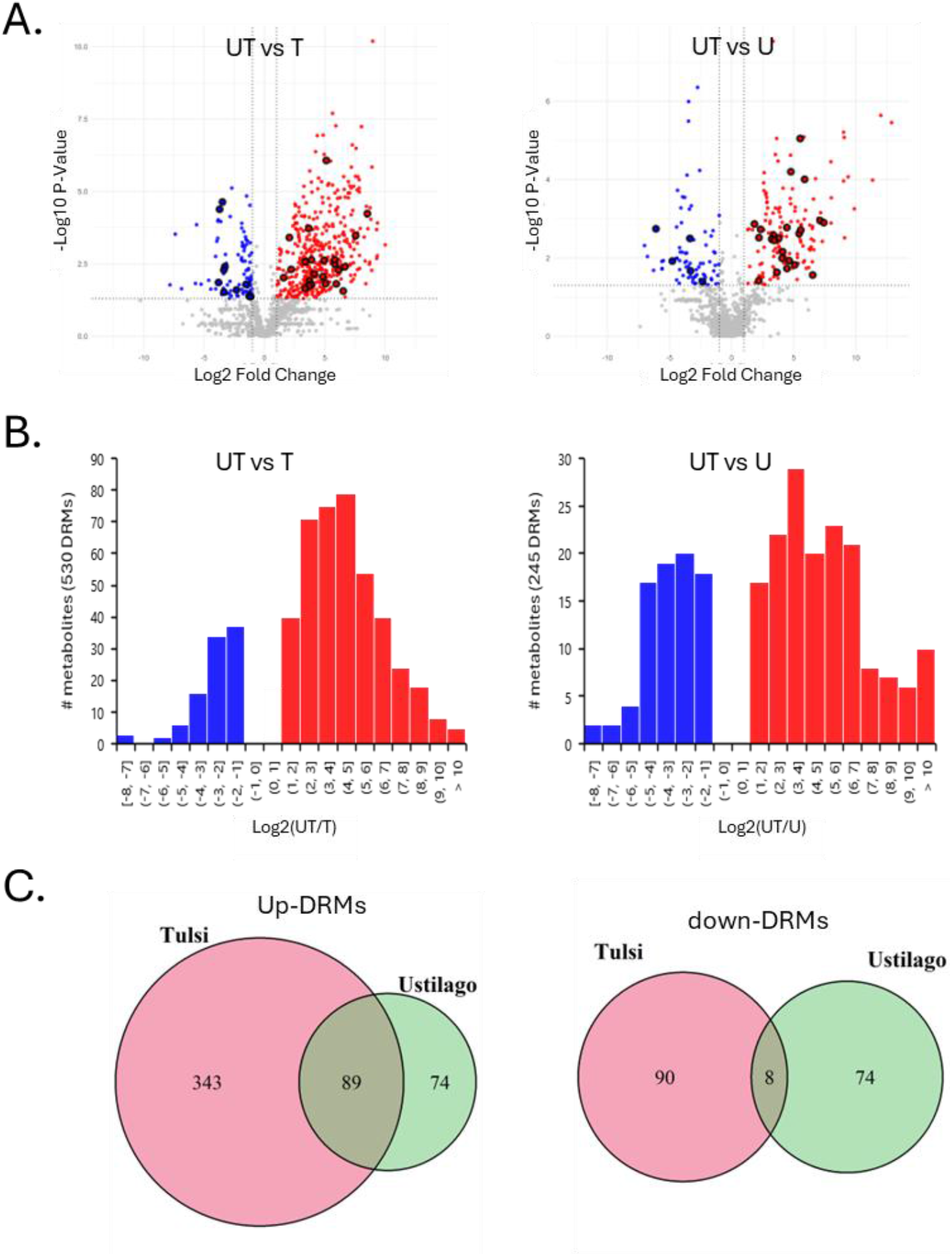
CFx-α1 enriched metabolome. A. Volcano plots of metabolites in Ustilago-Tulsi over the single Tulsi (left) or Ustilago (right). Up and down differentially regulated metabolites (DRMs) were highlighted with red and blue, respectively. Bolded circles mark antioxidant metabolites (see Figure 4A). B. Distribution of DRMs across Log2 fold change of LC-MS intensity in Ustilago-Tulsi over the single Tulsi (left) or Ustilago (right). C. Overlap between upregulated (left) and downregulated (right) DRMs shown in panels A-B. Mutually upregulated DRMs (89) are significantly enriched (p < 5.038e-13) by two-fold than randomly expected. U, T, and UT stand for Ustilago, Tulsi, and Ustilago-Tulsi (CFx-α1), respectively.

Specifically, the average log2 fold change of CFx-α1 up-DRMs versus T and U was 7.1 and 5.3, respectively, whereas the corresponding down-DRMs displayed average log2 fold changes of −3.4 and −1.7 (Figure 3B). Next, we compared the overlap between upregulated DRMs in CFx-α1 versus U and CFx-α1 versus T, as well as between downregulated DRMs (Figure 3C).Eighty-nine metabolites were found to be mutually upregulated in CFx-α1 relative to each of the single cultures, which is about twice the number expected by chance and statistically significant (p < 5.038 × 10^−13). In comparison, the overlap among mutually downregulated DRMs (n = 8) was enriched to a lesser extent (1.5-fold) and was not statistically significant (p > 0.05). No enrichment was observed between other DRM combinations, i.e., upregulated versus downregulated DRMs in either direction (Figure S2A–B). These results suggest that the CFx-α1 is enriched in metabolites compared with either of the single cultures and that the set of mutually upregulated metabolites is non-random.

### Synergistic upregulation of beneficial metabolites in the CFx-α1 metabolome

To further investigate the nature of mutually upregulated metabolites in CFx-α1 compared with single cultures, these metabolites were classified into four categories based on their presence in the single cultures: (1) metabolites present in both single cultures, (2) metabolites present in U but absent in T, (3) metabolites present in T but absent in U, and (4) metabolites absent from both U and T (Figure 4A). Metabolite intensity in CFx-α1 differed among the four groups, with the highest and lowest median intensities observed in the groups of metabolites present in both single cultures and absent from both single cultures, respectively (Figure 4B). The largest group with 40 mutually upregulated metabolites (45%) was the one with metabolites found in both single cultures (Figure 4A). Twenty-three metabolites, accounting for 26% of mutually upregulated metabolites, were found in only one of the single cultures (Figure 4A). Notably, a substantial fraction (29%) of the mutually upregulated metabolites in CFx-α1 belonged to the group that was absent from both single cultures (Figure 4A), suggesting de novo synthesis in the CFx-α1 metabolome. In addition, 85% of the mutually upregulated metabolites were enriched by at least fourfold in the CFx-α1 sample compared with the combined single cultures (U + T), implying a potent synergistic upregulation in CFx-α1 (Figure 4C). Vitamin C, polygodial, and octyl gallate are examples of metabolites from three different categories that were synergistically upregulated in CFx-α1 (Figure 4D). These metabolites have been reported to possess antioxidant, anti-inflammatory, antimicrobial, or wound-healing properties (32–34), suggesting potential health benefits of CFx-α1.

**Figure 4.**
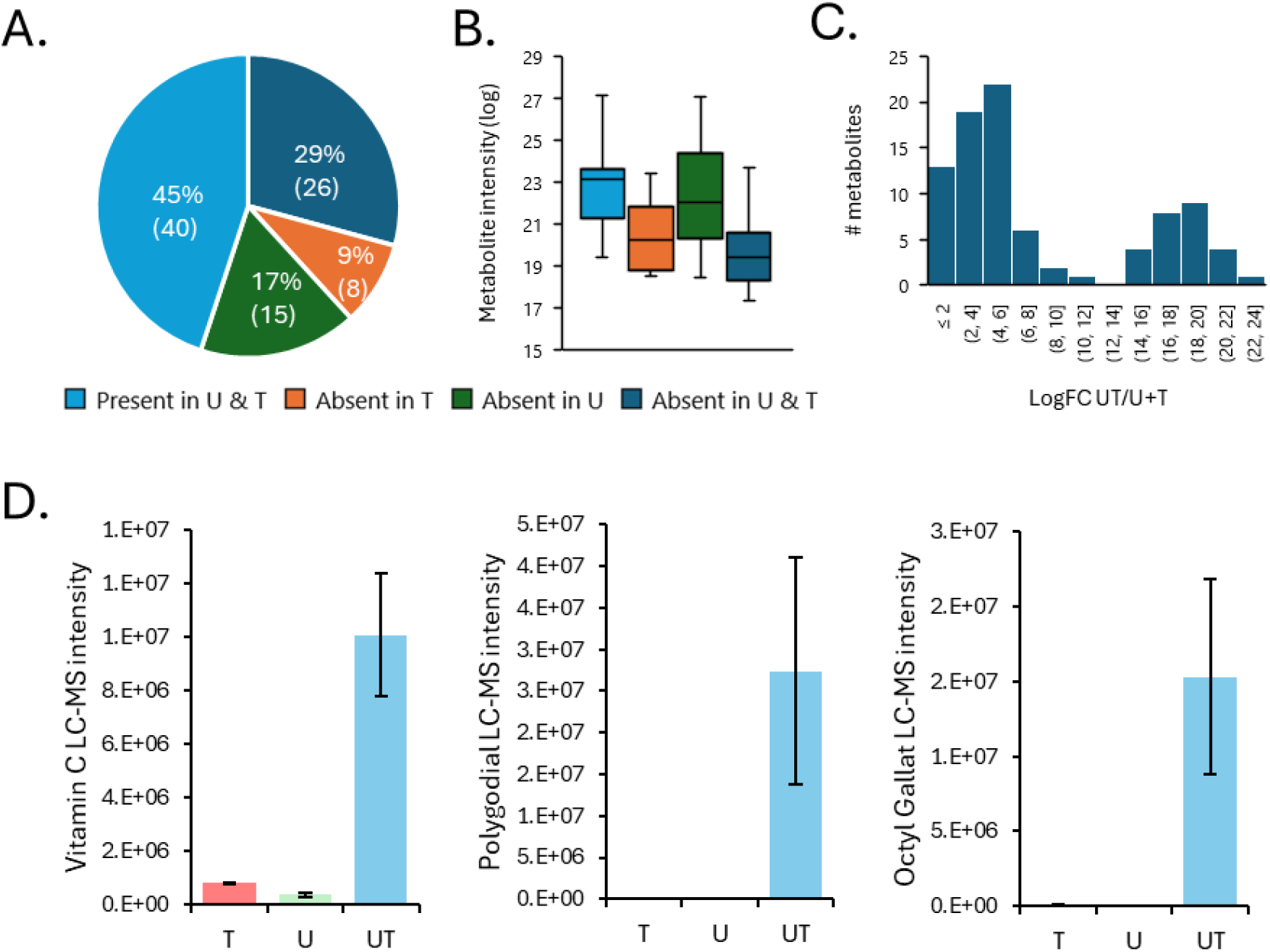
Synergistic upregulation of beneficial metabolites in the CFx-α1 metabolome. A. Proportions of the different categories of mutually upregulated DRMs in the Ustilago-Tulsi metabolome, i.e., that were upregulated in UT over U and T. B. Log intensity of metabolites in UT separated into four groups based on their presence status in the single cultures. C. Distribution of mutually upregulated DRMs in UT versus single cultures across Log2-fold change of metabolite intensity in UT versus the sum of single cultures. D. Metabolite LC-MS intensity of upregulated metabolites in UT that were present in U & T (Vitamin C), present only in one of the cultures (polygodial), or absent in both single cultures (octyl gallate). U, T, and UT stand for Ustilago, Tulsi, and Ustilago-Tulsi (CFx-α1), respectively.

### CFx-α1 is enriched in antioxidant metabolites and activity

Observing the activation of Vitamin C in the CFx-α1 metabolome, we next aimed to profile the general activation of antioxidant metabolites in each of the cultures. To do so, we developed an algorithm to annotate metabolites as antioxidants based on scientific publications (Figure 5A). The first step in the annotation process is searching for metabolites on PubMed. Here, we used the 1478 metabolites identified by the untargeted metabolomics data. Out of the total metabolites, 864 of them have been mentioned in papers, either in the title or abstract, along with the ‘antioxid*’ term. For accuracy, we excluded papers with partial match to our metabolite (i.e., modified metabolites), leaving us with 358 metabolites. In the next step, we ran the title and abstract through the GPT model to inquire about the direct role of the metabolite in antioxidation, which resulted in 246 metabolites. Manual reviewing suggested 95% accuracy of the GPT classification for metabolites with at least one positive/antioxidant classified paper (Figure 5B). The GPT success rate gradually increased with the number of positive classifications, reaching 100% with metabolites that at least five positive antioxidant classification, concluding 101 metabolites (Figure 5A).

**Figure 5.**
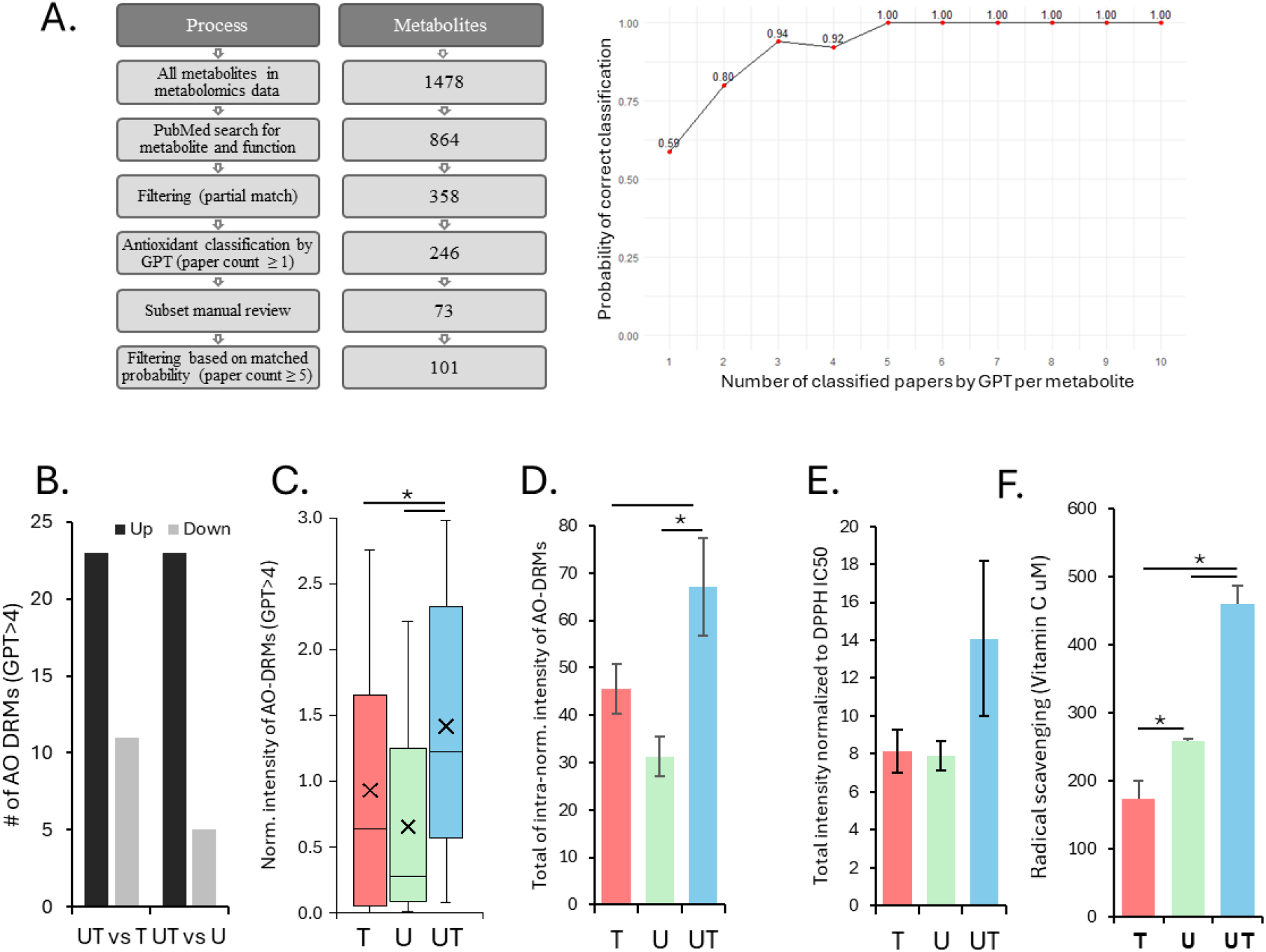
CFx-α1 is enriched in antioxidant metabolites and activity. A. A flowchart for the annotation of metabolites as antioxidants from scientific publications (left). Annotation accuracy of antioxidant increases with the number of positive publications (right). B. Number of up and down DRMs that were identified as antioxidants by our model (antioxidant scoring > 5 publications). C. Boxplot of normalized intensities (per-metabolite) of antioxidant metabolites. D. Combined intensity of antioxidant metabolites. Intensities were normalized per metabolite and metabolites. E. Total intensity of metabolites normalized intra-metabolite and to their DPPH IC50. F. Antioxidant activity based on the radical scavenging of DPPH. Antioxidant activity was normalized to Vitamin C DPPH scavenging. * marks significant differences (p<0.05). Values for all graphs is the average of three biological replicates. U, T, and UT stand for Ustilago, Tulsi, and Ustilago-Tulsi (CFx-α1), respectively.

Focusing on the 101 annotated antioxidant metabolites, we found that CFx-α1 to contain more antioxidants than either of the single cultures, i.e., 38 antioxidants compared to 32 for either U or T. Additionally, substantially more antioxidants were differentially upregulated than downregulated in CFx-α1 compared to either of the single cultures, specifically two times more in CFx-α1 versus U and five times more in CFx-α1 versus T (Figure 5B). Furthermore, quantification of LC-MS intensity showed antioxidants in CFx-α1 to be significantly more abundant than in the single cultures (Figure 5C-D). Antioxidants can exhibit a range of antioxidation potency. Therefore, we next normalized the abundance of antioxidants to their averaged published activity level and still found that the CFx-α1 samples had a higher total abundance of antioxidants than either of the single cultures (Figure 5E). Finally, we tested the antioxidant potency of the different samples using the DPPH radical scavenging assay (31). This analysis found that the CFx-α1 sample contained a radical scavenging activity equivalent to 460 mM Vitamin C, which was higher than that of either single cultures alone or combined (Figure 5F). Overall, these results suggest a robust and synergistic antioxidant potency of the CFx-α1 compared to the single cultures.

## Discussion

Our study demonstrates that yeast–plant-derived consortia factors (CFx-α1) possess a markedly enriched and distinct metabolomic profile compared with conventional cell-culture secretomes, underscoring the capacity of this biologically driven plant–microbial consortium to amplify beneficial bioactive compounds beyond what is achievable with standard cell culture systems. The metabolomic analysis revealed significant enrichment of antioxidants, organic acids, and bioactive metabolites in CFx-α1, suggesting synergistic metabolic interplay between the partners.

One particularly striking observation was the enhancement of vitamin C in the CFx-α1. While vitamin C was only mildly present in single cultures—approximately twice as high in Tulsi compared to Ustilago—its concentration increased dramatically in the CFx-α1, by 13-fold over Tulsi and 27-fold over Ustilago. This suggests that the consortium environment triggers metabolic pathways that neither organism activates alone. Possible mechanisms include cross-feeding of precursors, stress-induced signaling, and activation of host defense pathways, all of which are known to stimulate antioxidant biosynthesis(35,36).

Interestingly, this phenomenon parallels natural smut–host interactions. Huitlacoche, the edible corn smut caused by Ustilago maydis, has been reported to contain vitamin C among other nutrients (37). The enrichment of vitamin C in our CFx-α1 suggests that our cultivation process may mirror the natural interaction between smut fungi and their host plants, enabling similar metabolic enrichment. This finding underscores the ecological relevance of our approach and its potential for biotechnological applications.

Beyond vitamin C, the overall metabolomic profile of CFx-α1 indicates a broader enhancement of health-promoting compounds compared to single cultures. Such synergistic effects highlight the promise of consortium-based strategies for producing functional metabolites, offering a sustainable alternative to conventional single-organism fermentation systems(9,10).

Our findings—that repeat regions in *U. cynodontis* accumulate genetic variants at a rate (∼23/10 kb) nearly eight times higher than exonic regions (∼3/10 kb)—are consistent with trends across fungi and yeast, where repeat-rich compartments are hotspots for mutation while coding sequences are highly conserved (38). This supports the model of compartmentalized genome evolution, in which structural and regulatory diversity arises predominantly through repeat-associated variation, preserving essential protein-coding functions. In our strain, such dynamics may underpin subtle regulatory tuning that enhances secondary metabolism—such as elevated antioxidant production—without compromising vital genes.

The significant increase in vitamin C and other antioxidant and anti-inflammatory metabolites positions CFx-α1 as a powerful source of bioactive compounds with direct relevance to human health and wellness. Vitamin C is a cornerstone of immune support, collagen synthesis, and skin health, making these findings particularly impactful for the development of functional foods, nutraceuticals, and beauty formulations. By leveraging natural-like microbial–plant interactions, our approach offers a sustainable and scalable biotechnology platform to produce high-value compounds that promote skin and hair vitality, anti-aging benefits, and overall physiological resilience. These results open new avenues for utilizing cultivated Consortia Factors derived from plant and/or yeast cell cultures into health and beauty industries, bridging biotechnology with consumer well-being.

## References

1. Skalnikova H, Motlik J, Gadher SJ, Kovarova H. Mapping of the secretome of primary isolates of mammalian cells, stem cells and derived cell lines. Vol. 11, Proteomics. 2011.

2. Brown KJ, Seol H, Pillai DK, Sankoorikal BJ, Formolo CA, Mac J, et al. The human secretome atlas initiative: Implications in health and disease conditions. Biochim Biophys Acta Proteins Proteom. 2013;1834(11).

3. Caudy AA, Mülleder M, Ralser M. Metabolomics in yeast. Cold Spring Harb Protoc. 2017;2017(9).

4. Bhukya B, Banoth C, Keshav PK, Hussain MDS, Ajmera S. Biotechnological Production of Various Fungal Metabolites and their Applications in White Biotechnology. In 2022.

5. Wu T, Kerbler SM, Fernie AR, Zhang Y. Plant cell cultures as heterologous bio-factories for secondary metabolite production. Vol. 2, Plant Communications. 2021.

6. Bapat VA, Kavi Kishor PB, Jalaja N, Jain SM, Penna S. Plant Cell Cultures: Biofactories for the Production of Bioactive Compounds. Vol. 13, Agronomy. 2023.

7. Marchev A, Georgiev M. Plant Cell Bioprocesses. In: Current Developments in Biotechnology and Bioengineering: Bioprocesses, Bioreactors and Controls. 2017.

8. Sabater-Jara AB, Almagro L, Belchí-Navarro S, Martínez-Esteso MJ, Youssef SM, Casado-Vela J, et al. Suspension-cultured plant cells as a tool to analyze the extracellular proteome. Methods in Molecular Biology. 2014;1072.

9. Wijesekara T, Xu B. Health-Promoting Effects of Bioactive Compounds from Plant Endophytic Fungi. Vol. 9, Journal of Fungi. 2023.

10. Arora N, Patel A, Mehtani J, Pruthi PA, Pruthi V, Poluri KM. Co-culturing of oleaginous microalgae and yeast: paradigm shift towards enhanced lipid productivity. Vol. 26, Environmental Science and Pollution Research. 2019.

11. Bokelmann JM. Holy Basil/Tulsi (Ocimum tenuiflorum/sanctum). In: Medicinal Herbs in Primary Care. 2022.

12. Sen K, Goyal M, Mukopadayay S. review on phytochemical and pharmacological, medicinal properties of holy basil (Ocimum sanctum L.). Int J Health Sci (Qassim). 2022;

13. Digby S, Wells K. Compatibility and Development in Ustilago cynodontis. Mycologia. 1989;81(4).

14. Halisky PM, Webster RK. Heterothallism in Ustilago cynodontis (Pass.) Henn. Nature. 1963;197(4870).

15. Morita T, Konishi M, Fukuoka T, Imura T, Kitamoto D. Identification of Ustilago cynodontis as a new producer of glycolipid biosurfactants, mannosylerythritol lipids, based on ribosomal DNA sequences. J Oleo Sci. 2008;57(10).

16. Hosseinpour Tehrani H, Tharmasothirajan A, Track E, Blank LM, Wierckx N. Engineering the morphology and metabolism of pH tolerant Ustilago cynodontis for efficient itaconic acid production. Metab Eng. 2019;54.

17. Geiser E, Wiebach V, Wierckx N, Blank LM. Prospecting the biodiversity of the fungal family Ustilaginaceae for the production of value-added chemicals. Fungal Biol Biotechnol. 2014;1(1).

18. Newman N, Rajangam A, Talavera-Adame D, Sidhu H. System and method for the production, formulation and use of conditioned media, cultured cells and the factors included therein. USA: US 11,473,117 B2; 2019.

19. Ullmann L, Wibberg D, Busche T, Rückert C, Müsgens A, Kalinowski J, et al. Seventeen Ustilaginaceae High-Quality Genome Sequences Allow Phylogenomic Analysis and Provide Insights into Secondary Metabolite Synthesis. Journal of Fungi. 2022;8(3).

20. Li H, Durbin R. Fast and accurate short read alignment with Burrows-Wheeler transform. Bioinformatics. 2009;25(14).

21. Danecek P, Bonfield JK, Liddle J, Marshall J, Ohan V, Pollard MO, et al. Twelve years of SAMtools and BCFtools. Gigascience. 2021;10(2).

22. Talevich E, Shain AH, Botton T, Bastian BC. CNVkit: Genome-Wide Copy Number Detection and Visualization from Targeted DNA Sequencing. PLoS Comput Biol. 2016;12(4).

23. Stanke M, Diekhans M, Baertsch R, Haussler D. Using native and syntenically mapped cDNA alignments to improve de novo gene finding. Bioinformatics. 2008;24(5).

24. Stanke M, Schöffmann O, Morgenstern B, Waack S. Gene prediction in eukaryotes with a generalized hidden Markov model that uses hints from external sources. BMC Bioinformatics. 2006;7.

25. Cingolani P, Platts A, Wang LL, Coon M, Nguyen T, Wang L, et al. A program for annotating and predicting the effects of single nucleotide polymorphisms, SnpEff: SNPs in the genome of Drosophila melanogaster strain w1118; iso-2; iso-3. Fly (Austin). 2012;6(2).

26. Rempel A, Wittler R. SANS serif: Alignment-free, whole-genome-based phylogenetic reconstruction. Bioinformatics. 2021;37(24).

27. Bonini P, Kind T, Tsugawa H, Barupal DK, Fiehn O. Retip: Retention Time Prediction for Compound Annotation in Untargeted Metabolomics. Anal Chem. 2020;92(11).

28. Tsugawa H, Cajka T, Kind T, Ma Y, Higgins B, Ikeda K, et al. MS-DIAL: Data-independent MS/MS deconvolution for comprehensive metabolome analysis. Nat Methods. 2015;12(6).

29. Sumner LW, Amberg A, Barrett D, Beale MH, Beger R, Daykin CA, et al. Proposed minimum reporting standards for chemical analysis: Chemical Analysis Working Group (CAWG) Metabolomics Standards Initiative (MSI). Metabolomics. 2007;3(3).

30. Xia J, Psychogios N, Young N, Wishart DS. MetaboAnalyst: A web server for metabolomic data analysis and interpretation. Nucleic Acids Res. 2009;37(SUPPL. 2).

31. Blois MS. Antioxidant determinations by the use of a stable free radical [10]. Vol. 181, Nature. 1958.

32. Pullar JM, Carr AC, Vissers MCM. The roles of vitamin C in skin health. Vol. 9, Nutrients. 2017.

33. Kubo I, Fujita K, Lee SH. Antifungal mechanism of polygodial. J Agric Food Chem. 2001;49(3).

34. Park H, Ko R, Seo J, Ahn GY, Choi SW, Kwon M, et al. Octyl gallate has potent anti-inflammasome activity by directly binding to NLRP3 LRR domain. J Cell Physiol. 2024;239(4).

35. Shalaby S, Horwitz BA. Plant phenolic compounds and oxidative stress: integrated signals in fungal–plant interactions. Curr Genet. 2015;61(3).

36. Foyer CH, Noctor G. Redox homeostasis and antioxidant signaling: A metabolic interface between stress perception and physiological responses. Vol. 17, Plant Cell. 2005.

37. López-Martínez LX, Aguirre-Delgado A, Saenz-Hidalgo HK, Buenrostro-Figueroa JJ, García HS, Baeza-Jiménez R. Bioactive ingredients of huitlacoche (Ustilago maydis), a potential food raw material. Food Chemistry: Molecular Sciences. 2022;4.

38. Potgieter L, Feurtey A, Dutheil JY, Stukenbrock EH. On Variant Discovery in Genomes of Fungal Plant Pathogens. Front Microbiol. 2020;11.

