## Supplemental Figures for "Yeast–Plant-Derived Consortia Factors Exhibit A Distinct And Enriched Beneficial Metabolome For Health And Beauty"

A.

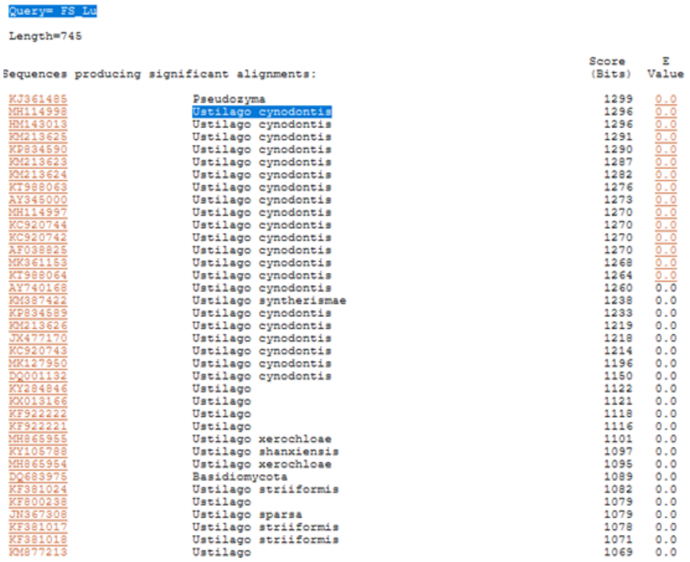


B.

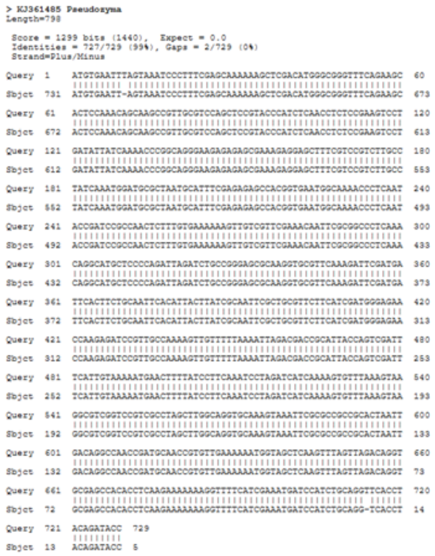


C.

| | SNP | Insertion | Deletion | Total Variants | Annotation length (bp) | Variants per 10kb |
|---|---|---|---|---|---|---|
| Exons | 3,585 | 162 | 180 | 3,927 | 11,823,229 | 3.3 |
| Introns | 263 | 24 | 34 | 321 | 522,471 | 6.1 |
| Intergenic | 3,790 | 558 | 506 | 4,854 | 7,178,230 | 6.8 |
| Repeats | 7,534 | 329 | 301 | 8,164 | 3,599,958 | 22.7 |
| Total | 15,172 | 1,073 | 1,021 | 16,694 | 23,130,474 | 7.2 |

**Supplementary Figure 1. Genetic profiling of the Ustilago culture**
A. Blast analysis of ITS sequence of isolated fungus culture. B. Homology of ITS sequencing between isolated fungus and U. cynodontis. C. Number and frequency of genetic variants in the genome of this study U. cynodontis strain vs the reference U. cynodontis NBRC 9727 strain.

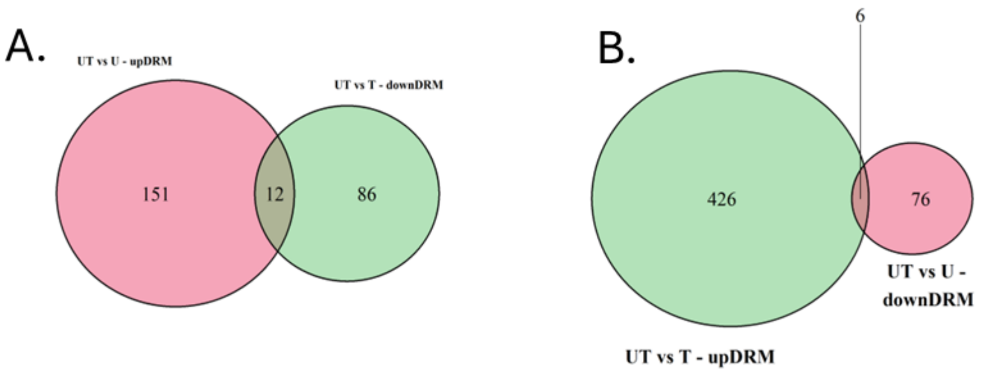


**Supplementary Figure 2. Overlaps between differentially regulated metabolites in UT vs U or T.**

A. Overlap between upregulated DRMs in UT vs. U (pink) and downregulated DRMs in UT vs T (green). B. Overlap between downregulated DRMs in UT vs. U (pink) and upregulated DRMs in UT vs T (green). U, T, and UT stand for Ustilago, Tulsi, and Ustilago-Tulsi (CFx-α1), respectively.
